# Effects of Reduced Achilles Subtendons Relative Displacement on Healthy Elderly Walking: A Simulation Study

**DOI:** 10.1101/2022.06.01.494269

**Authors:** Morteza Khosrotabar, Hamidreza Aftabi, Morad Karimpour, Majid Nili Ahmadabadi

## Abstract

Walking in healthy elderly people is characterized by lower performance. Since conventional training programs have had limited success in improving gait performance, it is essential to identify underlying causes of walking deficits in healthy elderly adults. Recent studies have qualitatively shown that the decreased relative displacement of Achilles subtendons is likely the primary contributor to lower propulsion in the elderly’s walking by creating a higher dependency on their triceps-surae muscle functions. Due to the invasive nature of experimental investigations, in this study, we developed a computational model and analyzed the effects of reduced Achilles subtendons relative displacement on the total metabolic rate and muscles’ force profiles during normal walking. Our musculoskeletal simulations revealed a 17% increase in the total metabolic rate in elderly adults whose Achilles subtendons were restricted to have no relative displacement. Changing the restriction level resulted in significant changes in the force distribution of the plantar flexor muscles, notably, a 40% reduction in the Medial Gastrocnemius and a 124% increase in the Soleus forces during the propulsion phase of walking. Also, we quantitatively presented the higher dependency of triceps-surae muscle functions regarding the limitation on their corresponding Achilles subtendons’ relative displacement. The results of this study confirm the experimental observations and can be used as initial insight into devising novel rehabilitation training programs with the focus on improving Achilles subtendons relative displacement.

## I. Introduction

**W**ALKING ability is one of the most important contributors to having a convenient daily life, which gradually degenerates by aging. According to the US Department of Health and Human Services, 17%, 28%, and 47% of elderly residents aged 65-74, 75-84, and over 85 years old, respectively, have walking difficulties which directly restrict their mobility [1]. In the last 20 years, researchers have conducted studies addressing the differences in walking characteristics between healthy young and older adults. Apart from time [2], length [3], and kinematic modifications [4], the most noticeable change from the older adults’ point of view is reduced propulsion during the push-off phase of walking [5]–[7], which results in a 0.2 to 0.8 m/s slower preferred speed [8], [9]. In this regard, it is well-documented that elderly people walk substantially with lower ankle moments and forward ground reaction forces compared with their younger counterparts [6], [10]. Compensating for their weaker propulsion, older adults utilize a distal to proximal shift mechanism in which they use hip and knee muscles more than healthy younger adults [7], [11]–[13], and due to the high energy consumption of those muscle groups, it is believed this compensatory mechanism is mainly responsible for the increased metabolic rate associated with elderly people [7], [14]. Identifying the underlying causes that lead to these defects has been a major concern of researchers for years to develop practical rehabilitation treatments.

Loss of muscle mass, which is usually considered to be a natural consequence of aging, is the first plausible explanation for the lower performance of elderly adults in walking. For a long time, researchers believed that the loss of triceps-surae muscle mass is the leading cause of diminished propulsion in elderly people and can be treated by resistance and power training programs [3]. However, there have been a lot of studies aimed at improving muscles’ strength through conventional training programs but did not find any positive correlation between the enhancement of muscles’ strength and improvements in propulsion metrics [15]–[18]. Moreover, researchers found that when older adults were asked to walk in conditions with higher external demands, such as uphill and fast walking, they utilized a reserve capacity in their muscles to complete the tasks [19]–[22]. Using the biofeedback, Brown *et al*. observed a substantial rise in the EMG activity of Gastrocnemius muscles and walking velocity in older adults when ask them to increase their forward ground reaction force and ankle power during the push-off phase of walking, respectively [23], [24]. Putting all these reports together, there is a consensus that loss of muscle mass is not sufficient to justify the reduced performance of elderly adults in walking. As the second vital component of humans’ propulsion system, the Achilles tendon is another potential defect source that needs to be explored.

The Achilles tendon is our strongest tendon and can bear forces almost four times the body weight [25], [26]. Like other body parts, the Achilles tendon goes through some changes in structure, shape, material composition, and mechanical properties with aging [27]–[29]. One of the significant changes is the lower stiffness of Achilles tendon in older adults compared with their younger counterparts [30], [31]. As the Achilles tendon becomes more compliant, it directly affects the operating length and velocity of the triceps-surae muscle fibers, changing their optimal functionality and force distributions [32]. Despite some efforts that attempted to increase the Achilles tendon’s stiffness using some training programs to prevent sports-related injuries [33]–[36], to the best of our knowledge, no study examined how these stiffening programs affected propulsion metrics in older adults.

Recent anatomical studies have revealed that the human Achilles tendon comprises three distinct subtendons, each of which is connected to one of the triceps-surae muscle bellies, the Soleus (Sol), the Medial and Lateral Gastrocnemius (MG & LG). By employing ultrasound imaging, researchers observed a non-uniform displacement between the deep (Sol subtendon) and superficial (MG and LG subtendons) layers of the Achilles tendon in various activities from level walking to passive ankle movements, and described this behavior as the Achilles subtendons relative displacement. It is believed this relative displacement is critical since it facilitates the independent function of our triceps-surae muscle [37], [38], and this becomes important when we know that our triceps-surae muscle groups play different roles in walking. Various experimental and simulation studies have tried to identify the unique function of Sol and Gastrocnemius muscles during walking cycles. Most of them have reported that the MG and LG muscles are the main contributor to forward propulsion, while Sol is responsible for body weight-supporting and helps Gastrocnemius muscles whenever the body encounters more demanding activities [39]–[41].

Ultrasound images of older adults during walking have revealed more uniform displacement profiles between the Achilles subtendons compared with young adults [42], [43]. In an animal study, Thorpe *et al*. saw a reduced relative displacement between the deep and superficial layers of the Achilles subtendons in older horses. They observed a higher stiffness within the interfascicular matrix of the Achilles tendon in aging horses, and due to the similarities between the humans’ and horses’ tendons, they believe that the same mechanism causes more uniform displacement across the humans’ Achilles tendon [44]. In support of this, Handsfield *et al*. simulated different interfascicular stiffness levels by considering various boundary conditions in a finite element model of the human Achilles subtendons. They found that higher stiffness values resulted in a lower relative displacement of the Achilles subtendons [45]. Decreased relative displacement creates a coupling among Sol, MG, and LG muscles and restricts them from performing at their optimal length and velocity. It is invasive to study the Achilles subtendons relative displacement through in-vivo experiments, thus one of the best ways to investigate this parameter is computational simulations. Accurate physiology-informed simulations in last decade provided the researchers with opportunities to study various aspects of human locomotion. In this study, we utilized musculoskeletal models from OpenSim and took advantage of Matlab software to investigate the effects of different relative displacement levels of the Achilles subtendons on muscles’ force distribution and total metabolic rate. For simplicity, we use subtendons sliding instead of the Achilles subtendons relative displacement in the rest of the text.

## II. Materials and Methods

### A. Experimental Data

We used the motion capture data of 26 lower limb markers and ground reaction forces of both legs collected and reported by Fukuchi *et al*. [46]. This well-gathered dataset comprises both overground and treadmill walking information at various speeds. In order to have a fair comparison between young and old groups and check the accuracy of our results, we chose seven subjects in each group from treadmill walking cases by considering three factors. First, the average speeds of elderly and young adults were close to each other (Old: 1.27 *±* 0.04 *m/s*, Young: 1.30 *±* 0.02 *m/s*); second, the selected speeds of individuals were the same or around their preferred speed, and third, we made the subject selection such that it maximises the age difference between elderly and young subjects (Old: 63.0 *±* 5.4 years, Young: 26.8 *±* 3.5 years). For transforming raw data (.c3d) to standard formats (.trc & .mot), we used the Mantoan *et al*. [47] converter toolbox.

### B. Calculating Muscles Force Distribution

First, we used Opensim software for scaling the generic musculoskeletal model *gait*2392 [48] to have similar anthropometry as our human subjects. We changed the upper body dimensions using the height of each subject combined with the lower limbs’ markers. Then, inverse kinematics and inverse dynamics techniques were utilized to find the joints’ angles and moments during one gait cycle (concerning heel strike of right foot).

In the next step, Matlab software was used for calculating the force distribution of muscles by solving the constrained optimization problem. Before defining optimization variables and constraints, constant properties of muscle-tendon units, like optimal fiber length, maximum velocity of muscle fiber, and tendon slack length, were imported from OpenSim’s generic model, and Thelen’s adjustments [49] were employed to modify these parameters for elderly adults.

We utilized the optimal control formulations of DeGroote *et al*. [50] for solving the muscle redundancy problem. In this approach, despite other time-marching solutions, dynamic equations are discretized and simulated by solving all time frames simultaneously. For solving the problem, the IPOPT [51] algorithm is called from the CasADi [52] toolbox in Matlab software. The cost function comprises of two terms. The first term represents muscular effort which is modeled by the sum of squared muscles excitations (*e*_*m*_), and the second term penalizes the usage of non-physiological reserve torque actuators:

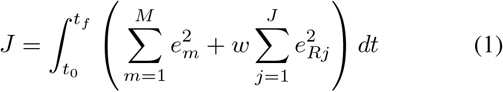

Where *t* is time, *t*_0_ and *t*_*f*_ are the initial and final time of the gait cycle, respectively, *m* = 1, …, *M* denotes the different muscles, *j* = 1, …, *J* indicates the different degrees of freedom (joints), and *e*_*Rj*_ is the input for *j*th reserve torque actuator.

Direct collocation method was selected over computed muscle control (CMC) algorithm [53], [54] for two main reasons. First, due to the combination of static optimization and forward simulation of muscle and skeleton dynamics, muscle forces computed with CMC are extremely sensitive to model parameter values [55]. More importantly, the direct collocation method gives us the ability to obtain the displacement of Achilles subtendons within triceps-surae muscle-tendon units. By having the subtendons displacement values, we can define our decreased subtendons sliding model.

### C. Decreased Subtendons Sliding (DSS) Model

Generally, in the DSS model we use the triceps-surae forces, which has already been calculated (Independent mode) to determine the maximum relative displacement between Achilles subtendons. Then, we impose a specific constraint on subtendons displacement and solve the optimization problem again to find the muscles force distribution considering the limitation of decreased subtendons sliding (Coupled modes). Here, for simplicity, we elaborate the DSS model between the Sol and MG subtendons, but in the main simulation it is applied on each two pairs of triceps-surae muscle-tendon units.

To put it more clear, as shown in Fig. 1, we assume a hypothetical situation where muscles and tendons are at their relaxed and rest state (slack length). We define points 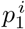 and 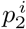 on Sol and MG subtendons, respectively, as corresponding points since they have the same distance from the calcaneus bone; i.e., they have the same rest lengths. In Fig. 1, a number of these corresponding points (blue on MG and green on Sol subtendon) are represented and connected by thin dashed lines. At an arbitrary time instance during the walking cycle (*t* = *t*_*k*_), when muscles are exerting forces, and subtendons are stretching, each pair of corresponding points move and create a relative displacement, 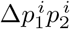. The closer the points are to the muscles, the greater their relative displacement is, thus the points 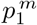 and 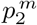 create the maximum relative displacement at each time instance: 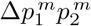. After calculating the muscles’ force distribution in the Independent Mode, we find the peak value of 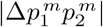 during the gait cycle for each subject and consider it as the subtendons sliding capacity of that person. To have a less sliding and lower relative displacement between subtendons in Coupled modes, we add DSS constraint (Eq. 2) to the optimization problem, and define the restriction level by adjusting the *λ* coefficient between zero and one.

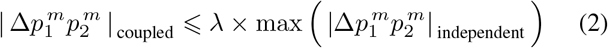

Considering *λ* = 1, does not impose a new limitation on the subtendons sliding, and the solver in this case finds the same solution as in the Independent mode. As *λ* decreases, the solver encounters a new situation and finds a different solution. At *λ* = 0, theoretically, there would be no sliding between Sol and MG subtendons. There are some studies in the literature that qualitatively discuss the relation between lower independent function of triceps-surae muscle-tendon units and lower sliding among their subtendons. In accordance with previous findings, we obtained a quantitative illustration depicting higher dependency among triceps-surae muscle functions.

**Fig. 1.**
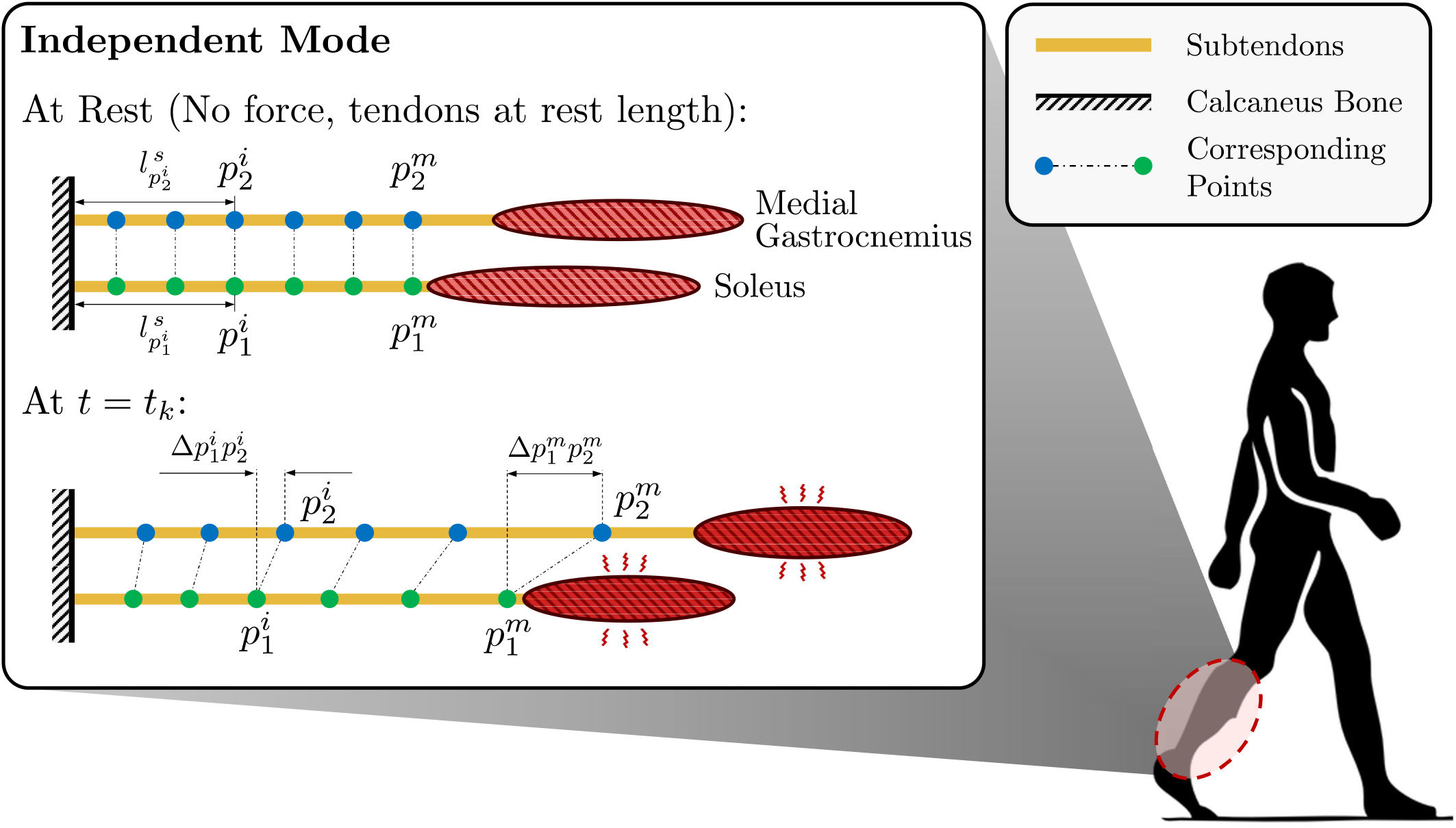
Decreased Subtendons Sliding Model. Sol and MG muscle-tendon units are shown in parallel. Corresponding points (blue and green points) have equal distances from calcaneus bone and are defined at the rest state, when there is no compression force in muscles and tendons are at their rest length. At *t* = *t*_*k*_, Sol and MG muscles are exerting forces and their subtendons are stretching, which creates a relative displacement between corresponding points, 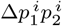. Further away the points are from the calcaneus bone, the greater the relative displacement is, thus 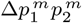 is the maximum relative displacement between the Sol and MG subtendons at each time instance.

### D. Higher Dependency among Triceps-Surae Functions

By utilizing the DeGroote muscle-tendon model [50] and considering zero sliding between the Achilles subtendons (*λ* = 0), we find an equation describing the relationship between triceps-surae muscle forces at all of the time instances during the gait cycle:

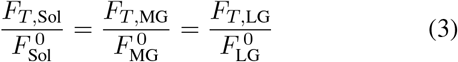

Where superscript 0 denotes isometric forces, and subscript *T* stands for the projected forces of muscles along with their subtendons (details are available in the Appendix section).

Pearson correlation of activation signals was used to show the linear relationship between triceps-surae muscle functions. As expected, and is shown in Fig. 2 the correlation strength increases significantly (*p <* 0.01) as *λ* decreases from Independent mode (*λ* = 1) to highly coupled mode (*λ* = 0.001).

**Fig. 2.**
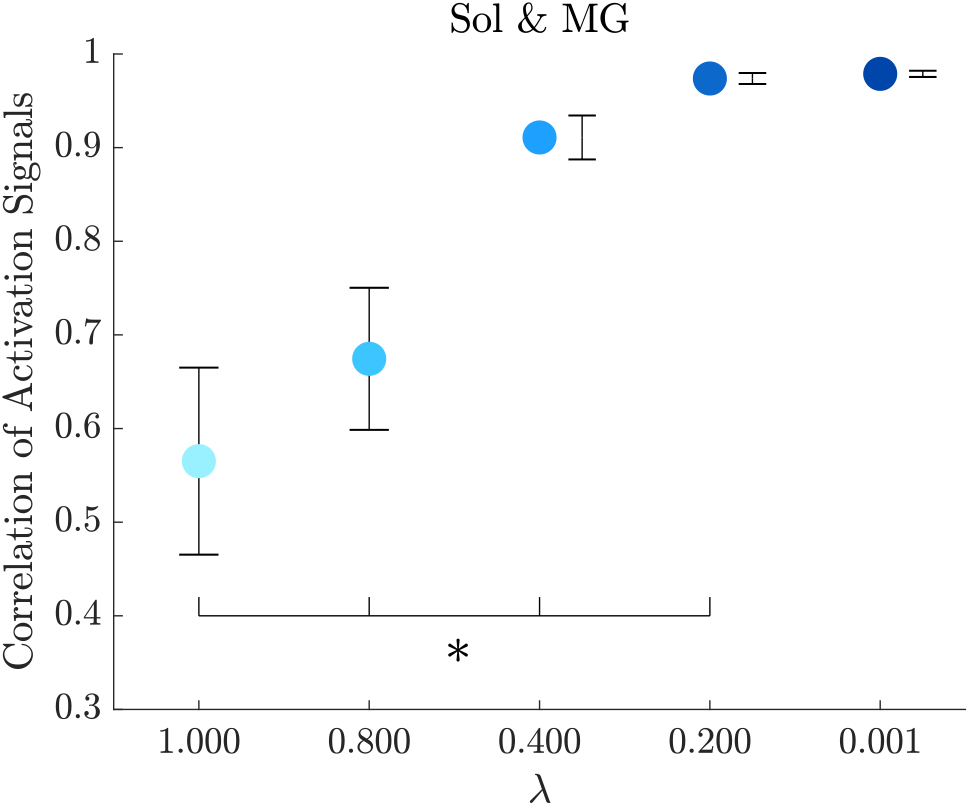
Correlation of activation signals between Soleus and Gastrocnemius muscles (Mean *±* SEM). Decreasing relative displacement of Achilles subtendons, increases the linear relation between Soleus and Gastrocnemius activation signals significantly (Two tailed paired t-test: *\*p <* 0.01).

### E. Statistical Analysis

In this study, a two-sided t-test with a significance level of 5% was used to show the significant effect of subtendons’ sliding on muscles’ force distribution during the propulsion phase of walking (30% to 60% of gait cycle) and total metabolic rate. To show the importance of subtendons sliding along with gait kinematics and generic model properties (old or young parameters), we utilized a regression model with F-statistic of 5% significance level.

### F. Assumptions, limitations, and considerations

#### 1) Same dataset in different λ values

The goal of this study is to investigate the effects of different *λ* values on the muscle’s behavior and total body metabolic rate, and since we have the same gait dataset in different *λ* values, it is not possible to assess other propulsion criteria like ankle moment or forward ground reaction force. Considering the DSS model within predictive simulations would be a good solution to find the effects of different subtendons sliding levels on such propulsion metrics.

#### 2) Modeling cause or effect

As mentioned before in the Introduction section, there are some evidence from animal studies that by getting older, the subtendons’ interfascicular stiffness increases and restricts their relative displacement [44]. The first approach toward simulating this behavior could be the cause-modeling. In this view, a material with variable stiffness is positioned between subtendons, and its stiffness level determines the subtendons relative displacement. Specifically, suppose the Sol and MG subtendons want to create the same relative displacement as the Independent Mode with stiffer interfascicular content. In that case, the Sol and MG muscles need to make a higher force difference than the Independent mode, and since this is not an optimal strategy, the body would prefer to lower the relative displacement of subtendons at higher stiffness levels. Another approach, the same as the one utilized in the DSS model, is to directly simulate what has been observed in the experimental studies, more uniform displacement or less sliding between subtendons. We applied the second approach here because, to the best of our knowledge, there were no human studies that approved the age-related increasing stiffness within the interfascicular matrix of subtendons. Furthermore, due to the complex geometry of subtendons during the gait cycle, it is not feasible to model a material with variable stiffness to properly restrict the subtendons from sliding.

## III. Results and Discussion

In this study, the effects of different subtendons sliding levels on muscles’ force distribution and total metabolic rate were investigated by considering four different *λ* values. Nine major lower body muscles were selected based on their isometric force and configuration (mono or bi-articular), and their responses to the DSS model were analyzed. Using the method presented by Umberger *et al*. [56], [57], the total metabolic rate was calculated for each subject. The specific tension of muscles and the ratio of fast-twitch fibers were reduced by 20 and 10%, respectively, to account for the age-related changes in the metabolic rate calculation [58].

### A. Muscles’ Force Distribution

Altering subtendons sliding directly affect muscles’ force distribution. Fig. 3 illustrates the average force profiles of nine major lower limb muscles at different *λ* values. As mentioned before, the gait kinematics and kinetics do not change at different *λ* values; therefore, we have the same torque profiles at each joint (inverse dynamics) that must be generated by muscles spanning that joint.

**Fig. 3.**
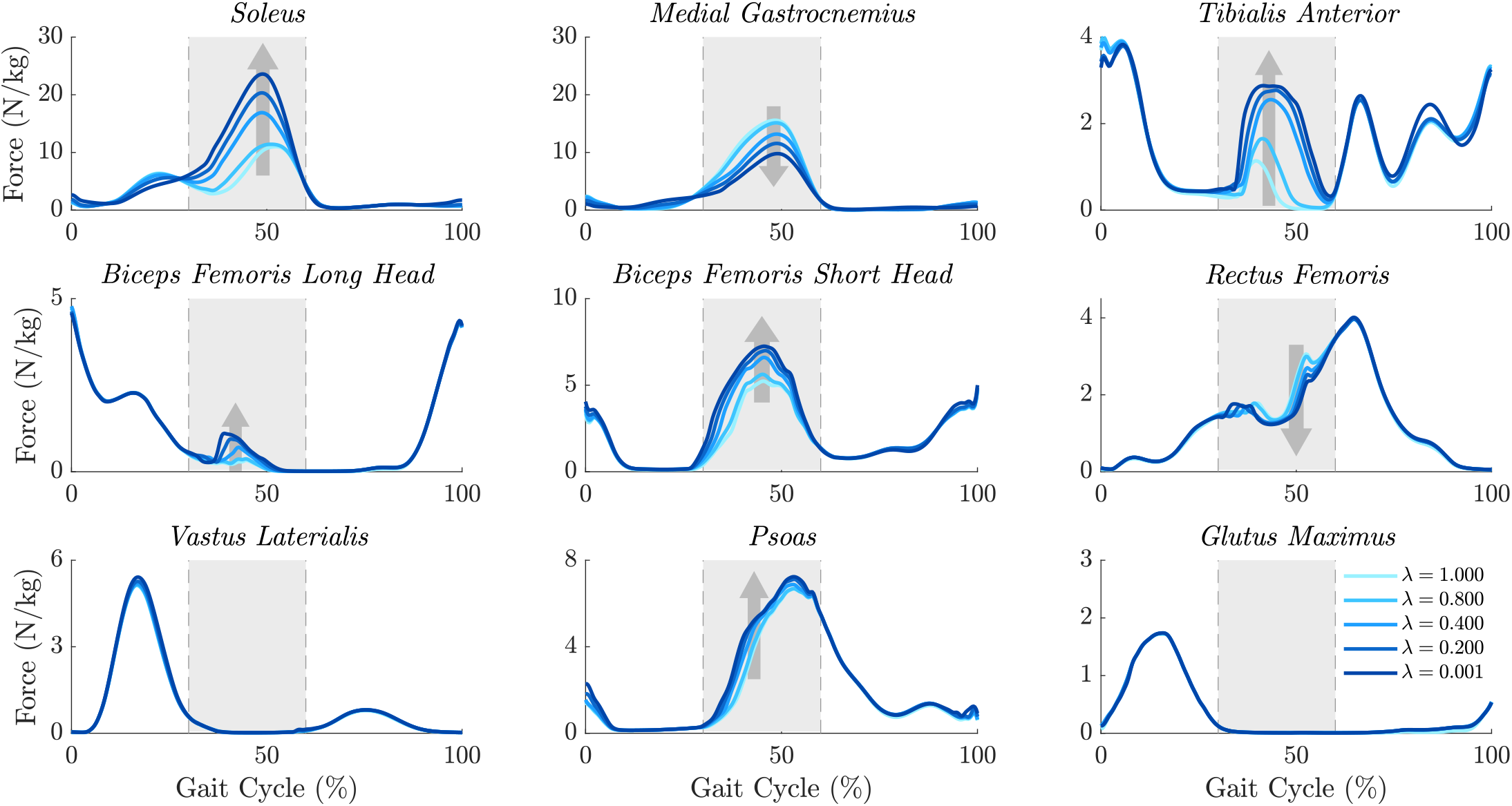
Muscles Force Distribution in different subtendons sliding levels. Lowering the relative displacement of Achilles subtendons results in a substantial changes of lower limb muscles force profiles during the propulsion phase of walking (30% to 60% of gait cycle) which is shown by the gray area. Nine major muscles regarding their isometric force and configuration (mono/bi-articular) are selected. *λ* = 1 indicates the Independent mode and as *λ* decreases, sliding between Achilles subtendons is restricted more intensely.

According to the linear relationship between Sol and MG forces (Eq. 3), if MG force increases or remains constant, we get much higher forces of Sol muscle during the propulsion phase, and to compensate for the excessive plantar-flexion torque at the ankle joint, the dorsiflexor muscles such as the Tibialis Anterior must increase their force substantially. The optimal approach would be having the MG force decreased, but it should be noted that the MG is a bi-articular muscle, and reducing its force to any extent can increase the activity of other knee flexors.

Our simulations with different *λ* values resulted in average 40% decrease (*−* 4.1 *N/kg*) of MG force during propulsion phase in older adults. As a result, Sol force increased by 124% (+8.2 *N/kg*) to compensate for the lack of plantar-flexion moment. Due to the linear relationship between Sol and MG muscles, excessive Sol forces are generated, which would be balanced by 1.4 *N/kg* increase in Tibialis Anterior force. As a bi-articular muscle, MG has a direct effect on knee flexion. Decreasing MG force leads to less knee flexion torque and must be compensated with higher activity of other knee flexor muscles as well as lower moments from knee extensor muscles. From the optimization point of view, the better approach would be adjusting the mono-articular muscles, as they do not affect other joints. Thus, as depicted in Fig. 3, Biceps Femoris Long/Short Head muscles increase their force by 0.25 *N/kg* and 1.67 *N/kg* respectively. Although decreasing the Vastus Laterialis force seems a sound solution due to its mono-articular configuration, as shown in Fig. 3, Vastus Laterialis force is zero during the propulsion phase of walking and cannot be reduced further, hence Rectus Femoris decreases its force slightly (11%,*−* 0.23 *N/kg*) to create the required torque at the knee joint. The same occurs at the hip joint and Psoas muscle increases its force by 19% (+0.76 *N/kg*) to meet the torque constraint from inverse dynamics.

An important interpretation of these changes could be the possibility of reversing the current paradigm by implementing an appropriate training program. As mentioned in the introduction section, Gastrocnemius muscles are believed to play a central role in forward propulsion during walking, and our simulations showed a substantial reduction in their force levels due to the lower subtendons sliding. Suppose the DSS model was true and humans have the exact mechanism toward aging. We can likely improve walking ability and especially propulsion intensities by designing training programs to facilitate the relative displacement of Achilles subtendons and enhance the independent function of triceps-surae muscle functions.

### B. Metabolic Rate

Having found the force distribution of lower limb muscles at different *λ* values, we calculated the total metabolic rate of elderly subjects, and as shown in Fig. 4, by decreasing the sub-tendons relative displacement, we observed a significant 17% increase (two tailed paired t-test, *p <* 0.01) in total metabolic rate on average. A significant greater contribution in total metabolic rate was observed from the muscles spanning the Knee (+54%, +0.45 *W/kg*) and Hip (+57%, +0.48 *W/kg*) joints compared with the Ankle joint (+28%, +0.23 *W/kg*). Based on these results, it seems that lower subtendons sliding in older adults might account for higher metabolic rates, and more importantly, improving subtendons relative displacement, may lead to a decreased total metabolic rate.

**Fig. 4.**
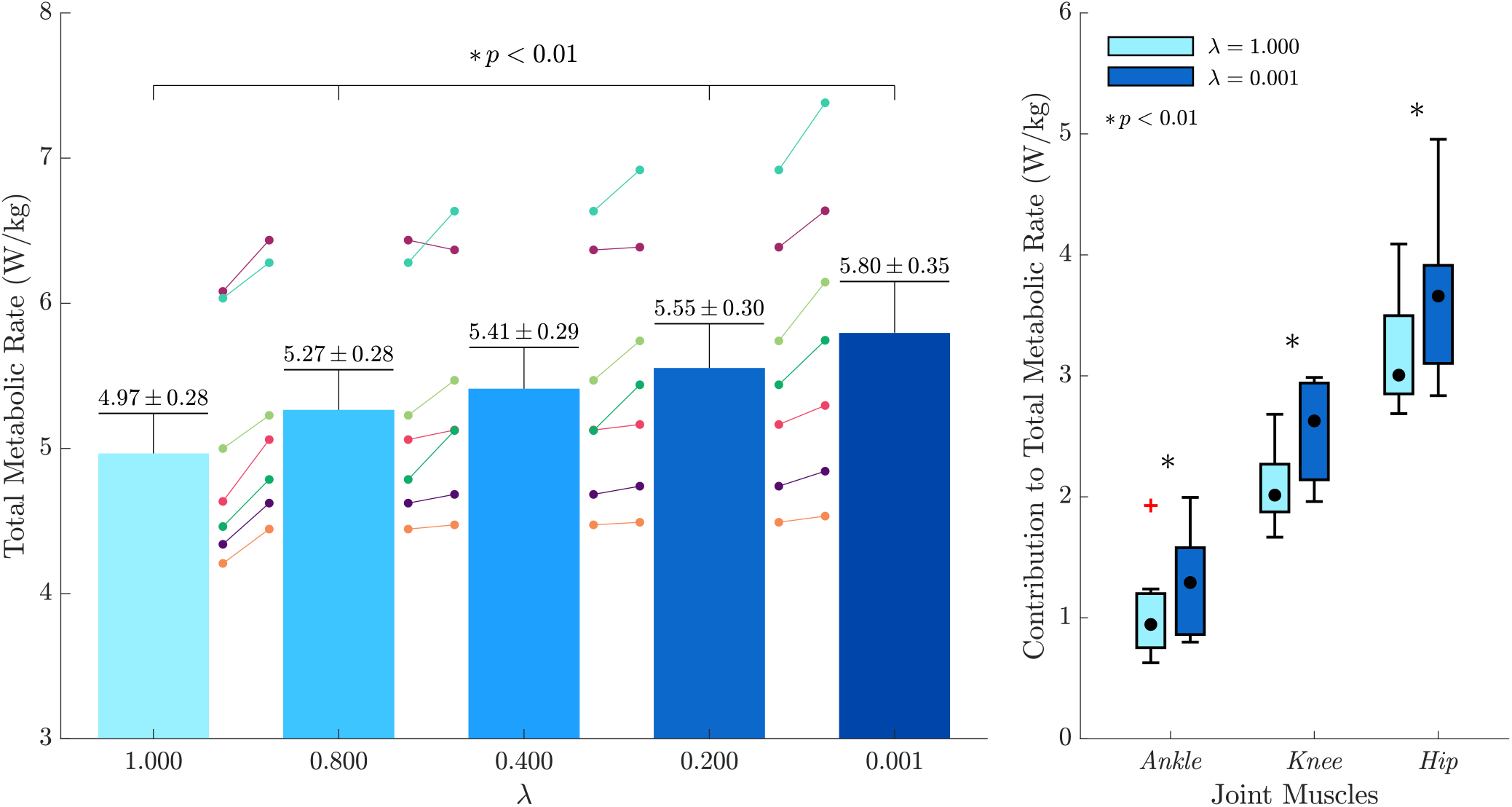
Effect of DSS Model on metabolic rate. Left: Decreasing the subtendons sliding results in a significant increase (+17%) of total metabolic rate in healthy elderly adults (Tow tailed paired t-test: *\* p <* 0.01). Right: Hip and knee muscles have a greater contribution to the higher total metabolic rate compared with the muscles spanning ankle joint (Red plus indicates an outlier data).

One of the parameters affecting an individual’s metabolic rate during walking, is their gait kinematics. In some studies, the higher metabolic rate of elderly adults is linked to the wider range of motion of their hip joint [7]. Second determining factor is the age difference, taken into account using musculoskeletal parameters in simulations. In this regard, some studies have indicated that older adults tend to rely more on their hip and knee joints to compensate for the loss of ankle strength, which leads to a higher metabolic rate. To find the importance of subtendons sliding levels in determining the total metabolic rate along with kinematics and age factors, we created a regression model considering two other imaginary conditions.

In *Old age − Young kinematics* condition, we utilized the motion capture and GRF dataset of young individuals, changed the parameters of their musculoskeletal models (similar to the old subjects) during the simulation and investigated their metabolic rate for different subtendons sliding levels. A possible interpretation of this condition would be to ask older adults to walk in another routine, through the use of biofeedback techniques. The second imaginary condition is *Young age − Old kinematics*, in which we used old group’s dataset and utilized the young musculoskeletal parameters. We consider this condition as old adults who trained themselves and improve their muscular capacities but try to walk with their habitual kinematics.

By combination of age and kinematics factors and employing different subtendons sliding levels, we came to 140 observations. After calculating metabolic rates at all of these cases, a linear regression model was trained in Matlab (R2019a) considering one discrete decision variable (subtendons sliding levels (*y*_1_), imported with *λ* values) and two binary decision variables (kinematic (*y*_2_) and age (*y*_3_), *Young* = 0, *Old* = 1):

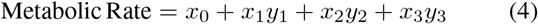

The results of this regression model are represented in Table I, and illustrate the importance of subtendons sliding factor for calculating the total metabolic rate. It seems that to have more reliable results in older adults, we should consider reduced subtendons relative displacement in musculoskeletal simulations. However, further clinical trials are needed to find the exact value of *λ* in each person and probably its changing rate by getting old.

**TABLE 1.** Results of regression model on total metabolic rates with 140 observations, rms-error = 0.536, r-squared = 0.535, and f-statistic vs. constant model = 52.2, p-value = 1.6e-22.

| Parameters | Coeff | Est | t-stat | p-value |
| --- | --- | --- | --- | --- |
| Intercept | $x_0$ | 4.28 | 43.70 | $6.1e-82$ |
| Subtendons Sliding ($\lambda$) | $x_1$ | -0.50 | -4.04 | $8.8e-05$ |
| Kinematics | $x_2$ | 0.32 | 3.54 | $5.4e-04$ |
| Age | $x_3$ | 1.02 | 11.30 | $2.6e-21$ |

## IV. Conclusion

In this paper, we developed a computational model to simulate reduced levels of Achilles subtendons sliding in elderly adults and investigated its effects on metabolic rate and muscles force distributions during the walking gait cycle. In particular, we utilized the musculoskeletal models from Open-Sim and solved the muscle redundancy problem by before and after applying a specific constraint on subtendons relative displacement. Our simulations revealed a significant increase in the total metabolic rate, along with considerable changes in muscles force distribution during the propulsion phase of walking. Furthermore, we found that the subtendons sliding levels influenced the metabolic rate as much as kinematics and age factors, and should be taken into account in musculoskeletal simulations. These findings can be used as a good insight into designing novel and appropriate training programs for older adults. In the case, by considering the effects of lower subtendons sliding, we can design a training cycle in which older adults increase their subtendons relative displacement and consequently improve their propulsion metrics.

## Appendix

### Higher Dependency of Triceps-Surae Muscle Functions

Having zero sliding (*λ* = 0) among the Achilles subtendons creates a linear relationship among triceps-surae muscle forces. Here we write the equations between Sol and MG, but it goes the same with LG muscle.

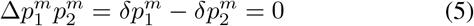

Where 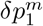 and 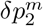 are defined using the nonlinear tendon model from DeGroote *et al*. [50]:

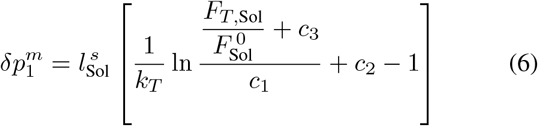

Details on tendon model and its constant parameters are available in [50]. 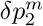 is defined as Eq. 6 except that MG is used instead of Sol within the bracket. By substituting the displacement of point 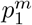 and 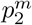 in Eq. 5, we have:

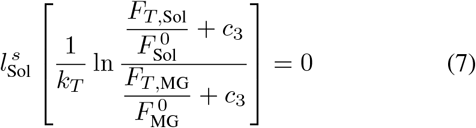

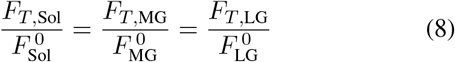

## References

[1] “ Physical Activity Guidelines Advisory Committee Scientific Report kernel description,” accessed: 2019. [Online]. Available: https://health.gov/sites/default/files/2019-09/PAGAdvisoryCommitteeReport.pdf

[2] D. A. Winter, A. E. Patla, J. S. Frank, and S. E. Walt, “ Biomechanical walking pattern changes in the fit and healthy elderly,” Physical therapy, vol. 70, no. 6, pp. 340–347, 1990.

[3] J. O. JudgeRoy, B. Davis III, and S. Õunpuu, “ Step length reductions in advanced age: the role of ankle and hip kinetics,” The Journals of Gerontology Series A: Biological Sciences and Medical Sciences, vol. 51, no. 6, pp. M303–M312, 1996.

[4] D. C. Kerrigan, L. W. Lee, J. J. Collins, P. O. Riley, and L. A. Lipsitz, “ Reduced hip extension during walking: healthy elderly and fallers versus young adults,” Archives of physical medicine and rehabilitation, vol. 82, no. 1, pp. 26–30, 2001.

[5] F. Prince, H. Corriveau, R. Hébert, and D. A. Winter, “ Gait in the elderly,” Gait & posture, vol. 5, no. 2, pp. 128–135, 1997.

[6] A. Silder, B. Heiderscheit, and D. G. Thelen, “ Active and passive contributions to joint kinetics during walking in older adults,” Journal of biomechanics, vol. 41, no. 7, pp. 1520–1527, 2008.

[7] J. R. Franz, “ The age-associated reduction in propulsive power generation in walking,” Exercise and sport sciences reviews, vol. 44, no. 4, pp. 129–136, 2016.

[8] J. E. Himann, D. A. Cunningham, P. A. Rechnitzer, and D. H. Paterson, “ Age-related changes in speed of walking.” Medicine and science in sports and exercise, vol. 20, no. 2, pp. 161–166, 1988.

[9] C. A. McGibbon and D. E. Krebs, “ Discriminating age and disability effects in locomotion: neuromuscular adaptations in musculoskeletal pathology,” Journal of applied physiology, vol. 96, no. 1, pp. 149–160, 2004.

[10] V. Monaco, L. A. Rinaldi, G. Macrì, and S. Micera, “ During walking elders increase efforts at proximal joints and keep low kinetics at the ankle,” Clinical biomechanics, vol. 24, no. 6, pp. 493–498, 2009.

[11] D. C. Kerrigan, M. K. Todd, U. Della Croce, L. A. Lipsitz, and J. J. Collins, “ Biomechanical gait alterations independent of speed in the healthy elderly: evidence for specific limiting impairments,” Archives of physical medicine and rehabilitation, vol. 79, no. 3, pp. 317–322, 1998.

[12] L.E. Cofré, N. Lythgo, D. Morgan, and M. P. Galea, “ Aging modifies joint power and work when gait speeds are matched,” Gait & posture, vol. 33, no. 3, pp. 484–489, 2011.

[13] P. DeVita and T. Hortobagyi, “ Age causes a redistribution of joint torques and powers during gait,” Journal of applied physiology, vol. 88, no. 5, pp. 1804–1811, 2000.

[14] T. Delabastita, E. Hollville, A. Catteau, P. Cortvriendt, F. De Groote, and B. Vanwanseele, “ Distal-to-proximal joint mechanics redistribution is a main contributor to reduced walking economy in older adults,” Scandinavian Journal of Medicine & Science in Sports, vol. 31, no. 5, pp. 1036–1047, 2021.

[15] P. Capodaglio, M. Capodaglio Edda, M. Facioli, and F. Saibene, “ Long-term strength training for community-dwelling people over 75: impact on muscle function, functional ability and life style,” European journal of applied physiology, vol. 100, no. 5, pp. 535–542, 2007.

[16] A. Hartmann, K. Murer, R. A. D. Bie, and E. D. D. Bruin, “ The effect of a foot gymnastic exercise programme on gait performance in older adults: a randomised controlled trial,” Disability and rehabilitation, vol. 31, no. 25, pp. 2101–2110, 2009.

[17] L. N. Persch, C. Ugrinowitsch, G. Pereira, and A. L. Rodacki, “ Strength training improves fall-related gait kinematics in the elderly: a randomized controlled trial,” Clinical Biomechanics, vol. 24, no. 10, pp. 819– 825, 2009.

[18] C. M. Beijersbergen, U. Granacher, A. A. Vandervoort, P. DeVita, and T. Hortobágyi, “ The biomechanical mechanism of how strength and power training improves walking speed in old adults remains unknown,” Ageing research reviews, vol. 12, no. 2, pp. 618–627, 2013.

[19] J. R. Franz and R. Kram, “ Advanced age affects the individual leg mechanics of level, uphill, and downhill walking,” Journal of biomechanics, vol. 46, no. 3, pp. 535–540, 2013.

[20] J. R. Franz and R. Kram, “ Advanced age and the mechanics of uphill walking: a joint-level, inverse dynamic analysis,” Gait & posture, vol. 39, no. 1, pp. 135–140,

[21] K. A. Conway and J. R. Franz, “ Increasing the propulsive demands of walking to their maximum elucidates functionally limiting impairments in older adult gait,” Journal of aging and physical activity, vol. 28, no. 1, pp. 1–8, 2020.

[22] K. A. Conway, K. L. Crudup, M. D. Lewek, and J. R. Franz, “ Effects of horizontal impeding force gait training on older adult push-off intensity.” Medicine and Science in Sports and Exercise, vol. 53, no. 3, pp. 574– 580, 2021.

[23] M. G. Browne and J. R. Franz, “ Does dynamic stability govern propulsive force generation in human walking?” Royal Society open science, vol. 4, no. 11, p. 171673, 2017.

[24] M. G. Browne and J. R. Franz, “ Ankle power biofeedback attenuates the distal-to-proximal redistribution in older adults,” Gait & posture, vol. 71, pp. 44–49, 2019.

[25] V. L. Giddings, G.S. Beaupré, R. T. Whalen, and D. R. Carter, “ Calcaneal loading during walking and running,” Medicine & Science in Sports & Exercise, vol. 32, no. 3, pp. 627–634, 2000.

[26] E. M. Keuler, I. F. Loegering, J. A. Martin, J. D. Roth, and D. G. Thelen, “ Shear wave predictions of achilles tendon loading during human walking,” Scientific Reports, vol. 9, no. 1, pp. 1–9, 2019.

[27] G. L. Onambele, M. V. Narici, and C. N. Maganaris, “ Calf muscletendon properties and postural balance in old age,” Journal of applied physiology, vol. 100, no. 6, pp. 2048–2056, 2006.

[28] L. Mademli and A. Arampatzis, “ Mechanical and morphological properties of the triceps surae muscle–tendon unit in old and young adults and their interaction with a submaximal fatiguing contraction,” Journal of Electromyography and Kinesiology, vol. 18, no. 1, pp. 89–98, 2008.

[29] L. Stenroth, J. Peltonen, N. J. Cronin, S. Sipilä, and T. Finni, “ Age-related differences in achilles tendon properties and triceps surae muscle architecture in vivo,” Journal of applied physiology, vol. 113, no. 10, pp. 1537–1544, 2012.

[30] T. Delabastita, S. Bogaerts, and B. Vanwanseele, “ Age-related changes in achilles tendon stiffness and impact on functional activities: a systematic review and meta-analysis,” Journal of Aging and Physical Activity, vol. 27, no. 1, pp. 116–127, 2018.

[31] B. K. Coombes, K. Tucker, F. Hug, and T. J. Dick, “ Age-related differences in gastrocnemii muscles and achilles tendon mechanical properties in vivo,” Journal of biomechanics, vol. 112, p. 110067, 2020.

[32] M. I. V. Orselli, J. R. Franz, and D. G. Thelen, “ The effects of achilles tendon compliance on triceps surae mechanics and energetics in walking,” Journal of biomechanics, vol. 60, pp. 227–231, 2017.

[33] A. Urlando and D. Hawkins, “ Achilles tendon adaptation during strength training in young adults.” Medicine and science in sports and exercise, vol. 39, no. 7, pp. 1147–1152, 2007.

[34] K. Kubo, T. Ikebukuro, H. Yata, N. Tsunoda, and H. Kanehisa, “ Time course of changes in muscle and tendon properties during strength training and detraining,” The Journal of Strength & Conditioning Research, vol. 24, no. 2, pp. 322–331, 2010.

[35] A. Arampatzis, A. Peper, S. Bierbaum, and K. Albracht, “ Plasticity of human achilles tendon mechanical and morphological properties in response to cyclic strain,” Journal of biomechanics, vol. 43, no. 16, pp. 3073–3079, 2010.

[36] J. M. Geremia, B. M. Baroni, M. F. Bobbert, R. R. Bini, F. J. Lanferdini, and M. A. Vaz, “ Effects of high loading by eccentric triceps surae training on achilles tendon properties in humans,” European Journal of Applied Physiology, vol. 118, no. 8, pp. 1725–1736, 2018.

[37] J. R. Franz, L. C. Slane, K. Rasske, and D. G. Thelen, “ Non-uniform in vivo deformations of the human achilles tendon during walking,” Gait & posture, vol. 41, no. 1, pp. 192–197, 2015.

[38] L. C. Slane and D. G. Thelen, “ Non-uniform displacements within the achilles tendon observed during passive and eccentric loading,” Journal of biomechanics, vol. 47, no. 12, pp. 2831–2835, 2014.

[39] R. G. Ellis, B. J. Sumner, and R. Kram, “ Muscle contributions to propulsion and braking during walking and running: insight from external force perturbations,” Gait & posture, vol. 40, no. 4, pp. 594–599, 2014.

[40] P. Malcolm, S. Galle, W. Derave, and D. De Clercq, “ Bi-articular kneeankle-foot exoskeleton produces higher metabolic cost reduction than weight-matched mono-articular exoskeleton,” Frontiers in neuroscience, vol. 12, p. 69, 2018.

[41] W. H. Clark, R. E. Pimentel, and J. R. Franz, “ Imaging and simulation of inter-muscular differences in triceps surae contributions to forward propulsion during walking,” Annals of Biomedical Engineering, vol. 49, no. 2, pp. 703–715, 2021.

[42] J. R. Franz and D. G. Thelen, “ Depth-dependent variations in achilles tendon deformations with age are associated with reduced plantarflexor performance during walking,” Journal of Applied Physiology, vol. 119, no. 3, pp. 242–249, 2015.

[43] J. R. Franz and D. G. Thelen, “ Imaging and simulation of achilles tendon dynamics: implications for walking performance in the elderly,” Journal of Biomechanics, vol. 49, no. 9, pp. 1403–1410, 2016.

[44] C. T. Thorpe, C. P. Udeze, H. L. Birch, P. D. Clegg, and H. R. Screen, “ Capacity for sliding between tendon fascicles decreases with ageing in injury prone equine tendons: a possible mechanism for age-related tendinopathy,” Eur Cell Mater, vol. 25, no. 4, 2013.

[45] G. G. Handsfield, J. M. Inouye, L. C. Slane, D. G. Thelen, G. W. Miller, and S. S. Blemker, “ A 3d model of the achilles tendon to determine the mechanisms underlying nonuniform tendon displacements,” Journal of biomechanics, vol. 51, pp. 17–25, 2017.

[46] C. A. Fukuchi, R. K. Fukuchi, and M. Duarte, “ A public dataset of overground and treadmill walking kinematics and kinetics in healthy individuals,” PeerJ, vol. 6, p. e4640, 2018.

[47] A. Mantoan, C. Pizzolato, M. Sartori, Z. Sawacha, C. Cobelli, and M. Reggiani, “ Motonms: A matlab toolbox to process motion data for neuromusculoskeletal modeling and simulation,” Source code for biology and medicine, vol. 10, no. 1, pp. 1–14, 2015.

[48] S. L. Delp, J. P. Loan, M. G. Hoy, F. E. Zajac, E. L. Topp, and J. M. Rosen, “ An interactive graphics-based model of the lower extremity to study orthopaedic surgical procedures,” IEEE Transactions on Biomedical engineering, vol. 37, no. 8, pp. 757–767, 1990.

[49] D. G. Thelen, “ Adjustment of muscle mechanics model parameters to simulate dynamic contractions in older adults,” J. Biomech. Eng., vol. 125, no. 1, pp. 70–77, 2003.

[50] F. De Groote, A. L. Kinney, A. V. Rao, and B. J. Fregly, “ Evaluation of direct collocation optimal control problem formulations for solving the muscle redundancy problem,” Annals of biomedical engineering, vol. 44, no. 10, pp. 2922–2936, 2016.

[51] A. Wächter and L. T. Biegler, “ On the implementation of an interiorpoint filter line-search algorithm for large-scale nonlinear programming,” Mathematical programming, vol. 106, no. 1, pp. 25–57, 2006.

[52] J. A. E. Andersson, J. Gillis, G. Horn, J. B. Rawlings, and M. Diehl, “ CasADi – A software framework for nonlinear optimization and optimal control,” Mathematical Programming Computation, vol. 11, no. 1, pp. 1–36, 2019.

[53] D. G. Thelen, F. C. Anderson, and S. L. Delp, “ Generating dynamic simulations of movement using computed muscle control,” Journal of biomechanics, vol. 36, no. 3, pp. 321–328, 2003.

[54] D. G. Thelen and F. C. Anderson, “ Using computed muscle control to generate forward dynamic simulations of human walking from experimental data,” Journal of biomechanics, vol. 39, no. 6, pp. 1107– 1115, 2006.

[55] M. Wesseling, F. De Groote, and I. Jonkers, “ The effect of perturbing body segment parameters on calculated joint moments and muscle forces during gait,” Journal of biomechanics, vol. 47, no. 2, pp. 596–601, 2014.

[56] B. R. Umberger, K. G. Gerritsen, and P. E. Martin, “ A model of human muscle energy expenditure,” Computer methods in biomechanics and biomedical engineering, vol. 6, no. 2, pp. 99–111, 2003.

[57] B. R. Umberger, “ Stance and swing phase costs in human walking,” Journal of the Royal Society Interface, vol. 7, no. 50, pp. 1329–1340, 2010.

[58] T. J. Doherty, A. A. Vandervoort, and W. F. Brown, “ Effects of ageing on the motor unit: a brief review,” Canadian journal of applied physiology, vol. 18, no. 4, pp. 331–358, 1993.

